# Cytoflow: A Python Toolbox for Flow Cytometry

**DOI:** 10.1101/2022.07.22.501078

**Authors:** Brian Teague

## Abstract

Cytoflow is a free, open-source flow cytometry toolbox that enables quantitative and reproducible analyses of flow cytometry experiments. Comprised of a set of well-documented Python modules wrapped by a graphical user interface, Cytoflow allows both programmers and bench scientists to apply modern data analysis methods (including machine learning) to high-dimensional flow data sets. Modern analyses may also lead to new insight about the biological systems that are studied with this powerful technique.

## Introduction

Flow cytometry is a technique for measuring the physical and chemical properties of individual cells as they flow through an instrument (1). In the most common instrument configuration (Fig 1), cells flow rapidly past one or several lasers, and the instrument measures scattered laser light as well as fluorescence in several channels. Scattered laser light gives insight into physical properties such as cell size and granularity, while fluorescent dyes, stains and proteins allow investigators to interrogate cellular properties and processes including cell viability (2), surface antigens (3), apoptosis (4), reporter gene expression (5), and many more (1).

**Figure 1.**
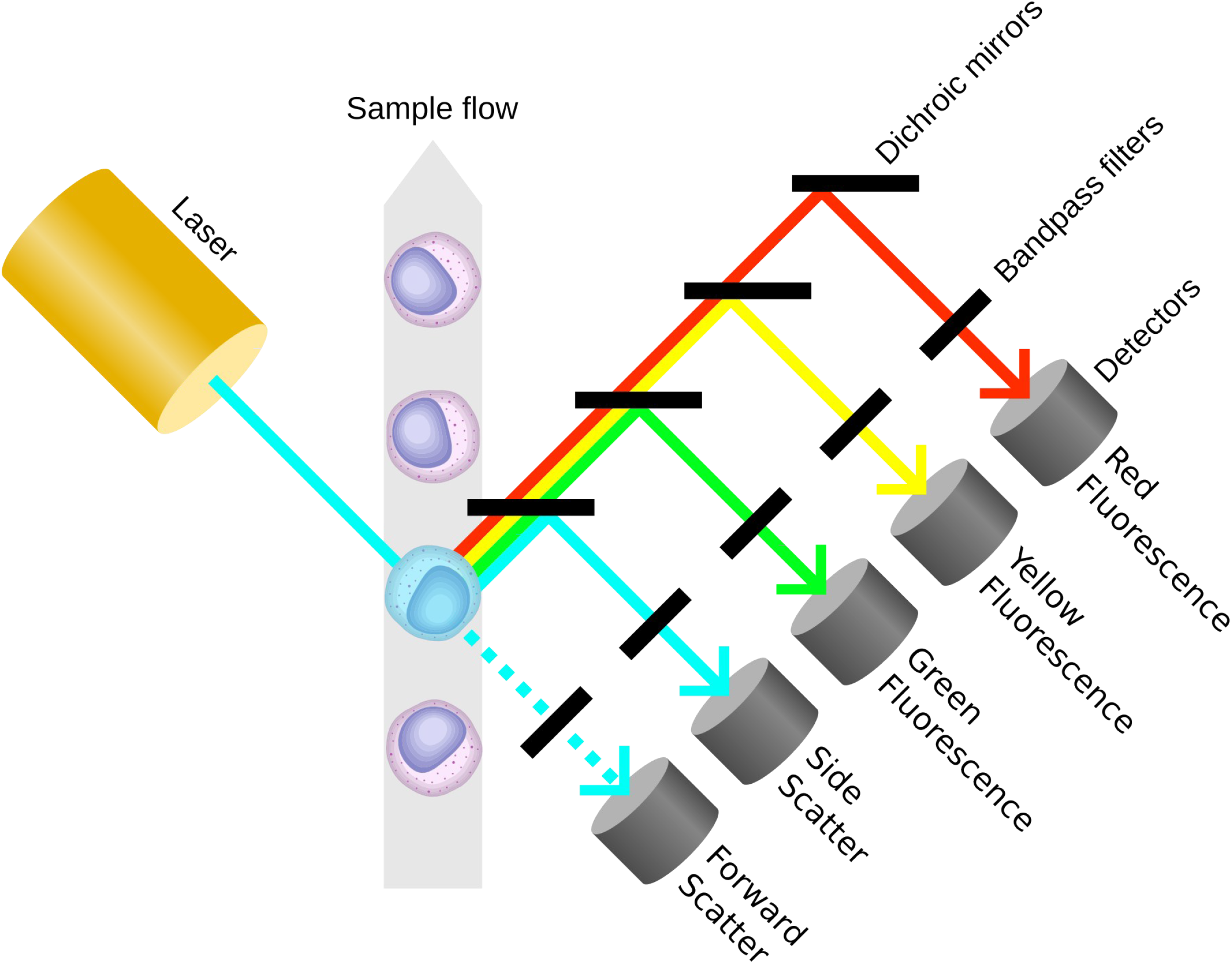
An optical flow cytometer analyzes cells’ fluorescence. Cells flow rapidly one-by-one past a laser, and a combination of mirrors, filters and detectors separate and quantitate both scattered laser light and fluorescence. Modern cytometers use multiple lasers and can measure more than 20 parameters from thousands of cells per second.

Because of its power and flexibility, many different experimental questions can be addressed with flow cytometry. Traditional cytometry often concerns itself with quantifying subpopulations in mixtures, counting the number of cells with a particular morphology or set of markers: is the cell a lymphocyte, monocyte, or neutrophil (6)? Are particular antigens present or absent? Is the cell alive or dead? However, as both instrumentation and reagents have improved, the types of problems addressed with flow cytometry have also begun to change. Modern experiments use the distributions of fluorescent signals – and their relationships – to build a quantitative, dynamical understanding of processes such as signal transduction and gene regulation (7,8), and current cytometry software often struggles to support these analyses. This is a particular issue in the field of synthetic biology, where distributions of single-cell fluorescence are a common way to characterize the performance of biological “parts” (9–11), but their performance may not be adequately captured in a simple statistic such as the cells’ mean fluorescence intensity (12).

Additionally, newer instruments produce data with higher dimensionality. Modern optical flow cytometers can analyze thousands of cells per second and report 20 or more parameters for each cell, resulting in very large multidimensional data sets (13). The field of data science continues to develop methods to gain insight from high-dimensional data (14,15), but these methods are only available to the scientists with the skills to manipulate these data programmatically.

Finally, this shift towards more sophisticated cytometry experiments has occurred alongside an increased emphasis on documented and reproducible analyses – a central tenet of the open science movement (16,17). Reproducibility is a particular problem for flow cytometry, because traditional analyses often rely on a skilled user to hand-draw “gates” – thresholds, ellipses or polygons – to identify populations of interest. Despite efforts by professional organizations (18), the positions of these gates are often not recorded and are almost never reported in publications. Thus, these analyses are often difficult to reproduce, and may also introduce unintentional biases into the conclusions (19).

Cytoflow is free, open-source flow cytometry software that addresses these three issues. Comprised of a set of reusable Python modules wrapped by a graphical user interface, Cytoflow allows both programmers and non-programmers to analyze flow cytometry data using approaches that are modern, quantitative and reproducible. Data can be transformed using traditional spectral compensation (20) or modern standards-based absolute quantification (21); classified using both hand-drawn gates and unsupervised machine-learning algorithms (22,23); and plotted using traditional histograms and scatterplots as well as modern multidimensional visualizations (24,25). The underlying Python modules are extensively documented and useful both in interactive data exploration as well as scripted analyses. Modern open-source development processes welcome contributions, patches, bug reports, and even whole-sale forking to adapt Cytoflow to new problems. Below, I describe Cytoflow’s design goals and implementation, give a detailed example of its use, and briefly touch on the project’s future directions.

## Design and Implementation

Cytoflow’s implementation addresses several design goals:

### Reproducible analysis

Whether explicit or not, most cytometry analyses proceed according to a “workflow,” a set of manipulations or transformations that are applied in order. For example, a cytometrist may open their data files (data loading), compensate the data for spectral overlap between adjacent channels (data transformation), view a plot of their data (data visualization), draw a gate to identify a subpopulation of interest (data selection), and then count the number of events in that gate (data reduction). Cytoflow makes this paradigm explicit (Fig 2): It encapsulates the data in an Experiment object, which contains the flow cytometry data as well as any statistics and metadata (discussed below). Data manipulations are implemented in Operation classes and visualizations in View classes (Fig 3). All Operations act as transformations on Experiments - they consume an Experiment and produce an Experiment. A set of fully-parameterized Operation instances, and the order in which they are applied to the raw cytometry data, constitute an analysis workflow that can be applied to new data, shared with collaborators, and reported upon publication.

**Figure 2.**
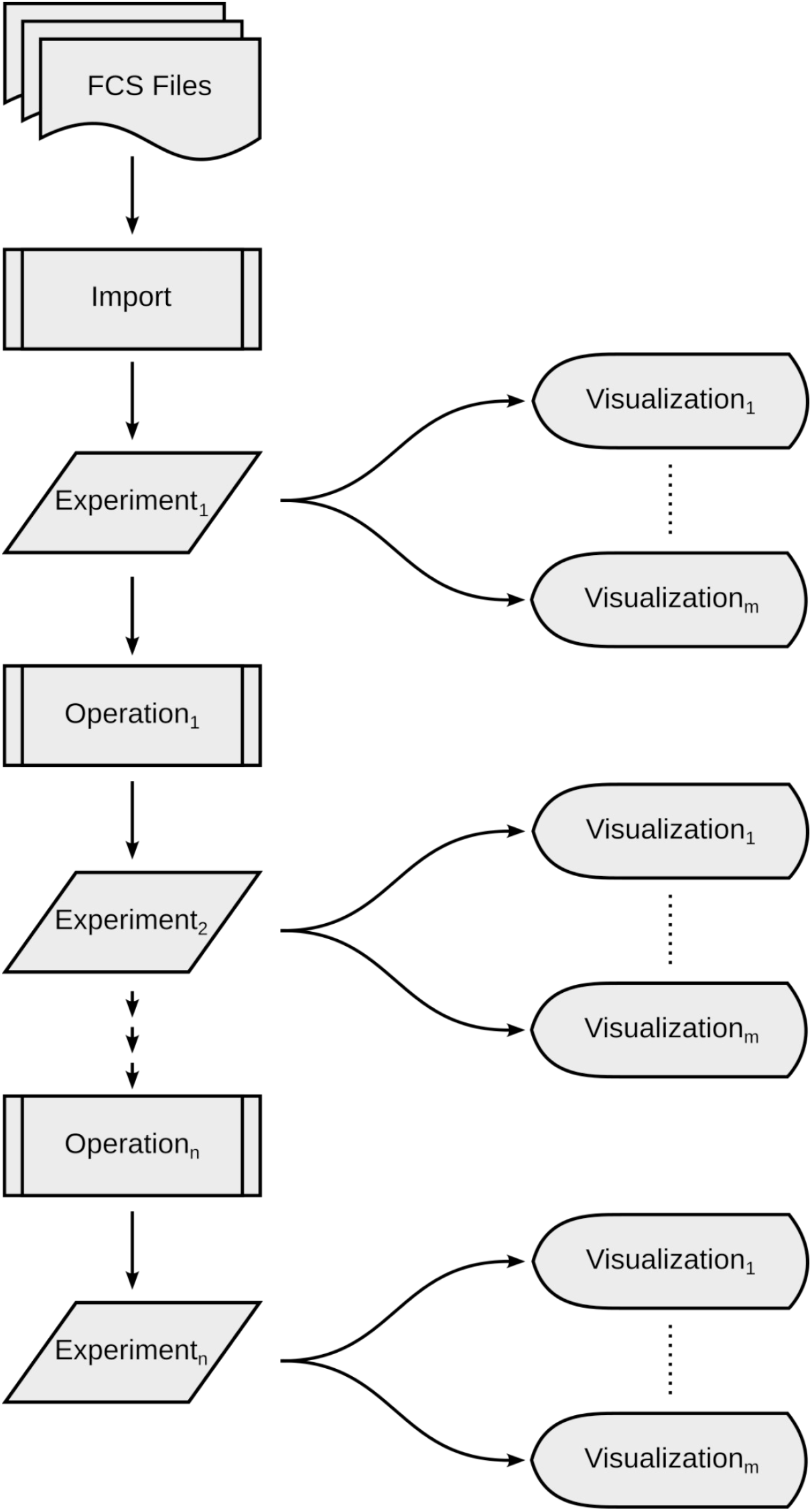
Flow cytometry data is analyzed with a “workflow”. FCS files are imported to create an Experiment, which is transformed by subsequence Operations. At any point in the workflow, Experiments can be visualized.

**Figure 3.**
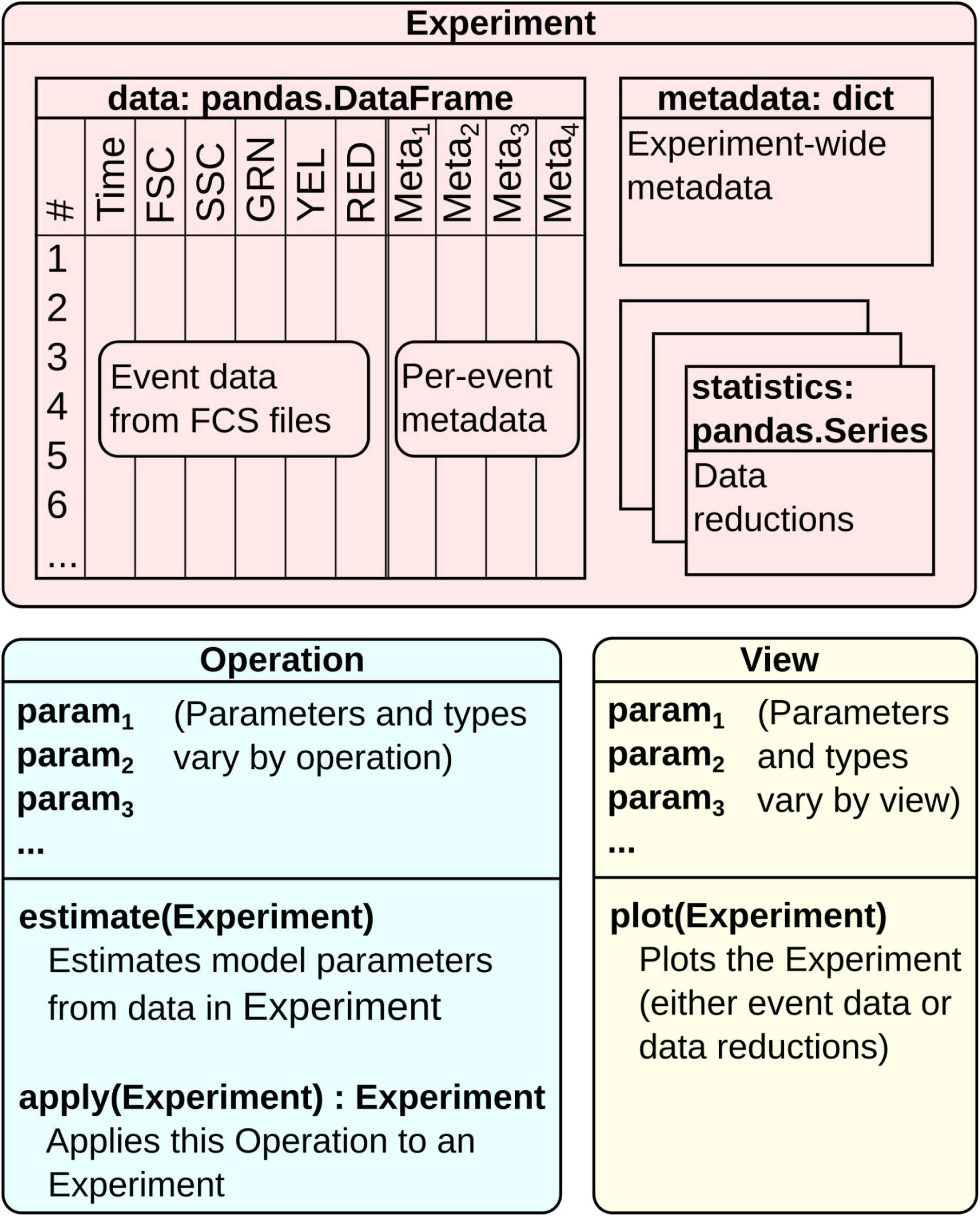
Cytoflow’s most important classes are Experiments, Operations and Views. An Experiment is Cytoflow’s primary data container – it holds event data from the FCS files, per-event metadata, per-experiment metadata and data reductions (statistics). Operations represent parametric data manipulations such as gating and spectral compensation. Some Operations’ parameters can be estimated from Experiments; others are set manually. Finally, Views are visualizations of flow cytometry data. They range from traditional histograms and scatterplots to sophisticated multidimensional representations.

### Intelligent metadata

In a research setting, flow cytometry experiments usually have experimenter-controlled independent variables: the concentration of a small-molecule inducer, or the genotype of the cells, or the treatment that an animal (or patient) received. Cytoflow uses these experimental metadata to structure its analysis and visualization, for example by plotting different experimental variables in different colors or grouping them before applying a data reduction. This is particularly powerful when analyzing experiments with multiple independent variables - no more trying to keep track of which tube or well had which conditions! Additionally, many Operations add per-event metadata to an Experiment which can be used in similar ways. For example, a Threshold operation adds a column of boolean metadata to the experiment which is true if the event is above the threshold and false otherwise. Per-event metadata keeps the entire dataset in a single “tidy” data structure (26) and allows subsequent Operations and Views to select and manipulate groups of data – for example, to color events on a plot or exclude subsets from further analysis.

### Powerful data reductions

A crucial step in most cytometry analyses is a data reduction that summarizes the raw events. Often, this is simply counting the number of events in a population of interest or determining its mean fluorescence in a particular channel. Cytoflow generalizes this process by allowing Operations to produce Statistics – quantities that are computed from the raw or transformed data. For example, an experimenter may want to know how the proportion of events above a threshold gate changes as their experimental variable changes – Cytoflow allows them to call length() (or any other valid Python function) on the subsets specified by per-event metadata, and the results are stored as a Statistic in the Experiment alongside the raw event data. Statistics are key objects in Cytoflow, which also provides several tools to visualize and manipulate them. In addition to using arbitrary Python functions to reduce and summarize data, a number of operations also produce Statistics as summaries of the data they are operating on, which are described in more detail below.

### Modern multivariate analyses and visualizations

Modern flow cytometers produce large, multivariate data sets, and Cytoflow gives users access to sophisticated analyses and visualizations to understand their data. For example, populations of interest are often “clusters” in a multidimensional space – Cytoflow includes several unsupervised machinelearning models, including k-means and Gaussian mixture models, to identify these clusters and assign events to them. These Operations also produce Statistics summarizing the model parameters, which are often useful data reductions themselves. For example, if the user fits Gaussian mixture models to various subsets of their data, the operation both assigns each event to a component and produces statistics about the components themselves, including their center, spread and the proportion of events that were assigned to them.

Cytoflow also makes multivariate visualizations easy. “Small multiples” (27) plots compare distributions across data subsets, and a number of modern visualizations such as RadViz(25) and parallel coordinates(24) are also included.

### Usable by both bench and data scientists

Cytoflow enables modern, reproducible flow cytometry analysis by both programmers and nonprogrammers. For data scientists, Cytoflow is available as both an Anaconda package (https://anaconda.org) and a Python wheel (https://pypi.org) and it integrates gracefully with data modern analysis environments such as the Jupyter Notebook (https://jupyter.org/), allowing for both data exploration and scriptable analyses. For non-programmers, the underlying Python modules are wrapped in a graphical user interface (GUI) that formalizes the “workflow” paradigm (Fig 4) while allowing access to all of Cytoflow’s operations and visualizations. Analyses that are created in the GUI can be saved without their data and applied to new data sets or exported as a Jupyter notebook. The GUI is supported on Linux, Mac and Windows systems and does not require installing a Python development environment.

**Figure 4.**
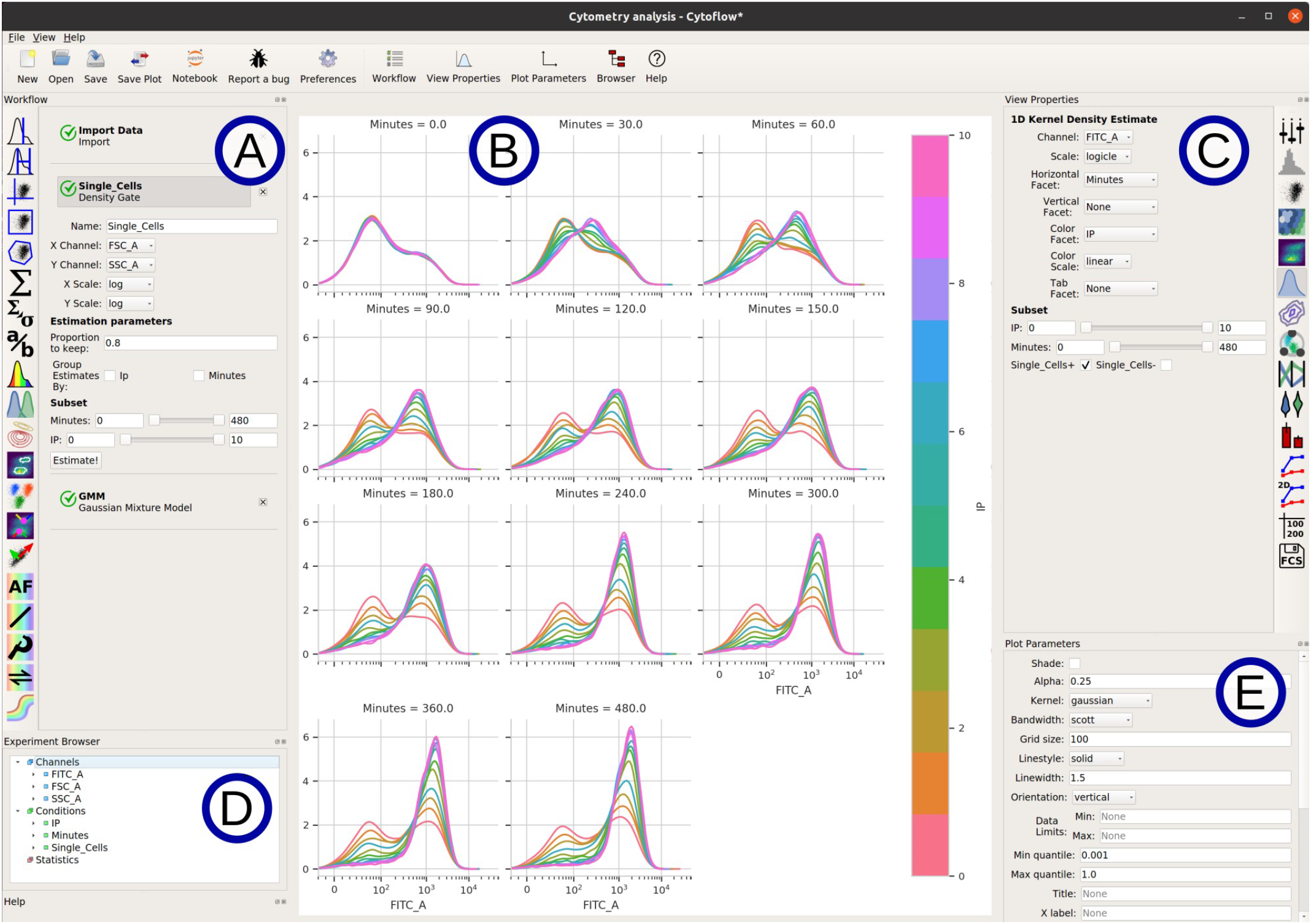
Cytoflow’s GUI wraps the underlying Python modules. (A) The Operations in the workflow are applied in order, beginning with an “Import Data” operation. Each Operation’s parameters can be set (or estimated) here as well. (B) The plot of the currently selected View. (C) The parameters of the currently selected view. (D) A browser for the current Experiment, including instrument channels, experimental variables, and statistics. (E) Additional plot parameters that are shared across views, including axis titles and limits.

### Easy to adopt and adapt

Cytoflow is an open-source project, allowing the underlying code to be inspected and modified by anyone with the requisite experience. Adaptation to new applications is easy because Cytoflow already uses common data structures and algorithms from the scientific Python ecosystem, and adding new Operations and Views is straightforward because the interface for each is only a few methods. Cytoflow is also extensively documented, with tutorials, HOWTOs and reference materials available for both the graphical user interface and the underlying Python modules (https://cytoflow.readthedocs.io). The project is hosted on GitHub, which centralizes development and community interaction.

## Results

The following analysis characterizes the time-varying response of a GFP-expressing yeast strain to varying concentrations of a small-molecule inducer. I induced several genetically engineered *Saccharomyces cerevisiae* cultures (28) with 12 different amounts of the small molecule *isopentyladenine* (IP) and measured the green fluorescence of the cultures on the cytometer over the course of the day, every 30 minutes for 8 hours. Full details, including both a step-by-step tutorial (demonstrating the GUI) and a runnable Jupyter notebook (demonstrating the underlying Python modules) can be found in the Tutorials section of the documentation (https://cytoflow.readthedocs.io), under ‘Yeast Induction Timecourse’.

### Step 1: Load the data and assign experimental variables

Cytoflow assumes that each FCS file corresponds to one tube or well of cells and requires that each be labeled with a unique set of experimental metadata (such as experimental conditions or replicate number). Thus, loading the data involves specifying the independent variables and assigning a value for each variable to each tube – in this case, “IP” (isopentyladenine concentration) and “Minutes” (the time that the sample was measured).

### Step 2: Select the single-cell population for analysis

Flow cytometers do not distinguish between single cells, cell fragments, and clumps of cells. Thus, most cytometrists use the forward-scatter and side-scatter information to select events that represent single cells and focus their analysis on these events. In this analysis, I applied a “density gate” (12) – it selects the largest contiguous area that contains a specified proportion of the events (Fig 5a). This will exclude cell fragments and clumps, which are often much smaller populations. Using a density gate is much less prone to operator error or bias than drawing a gate by hand.

**Figure 5.**
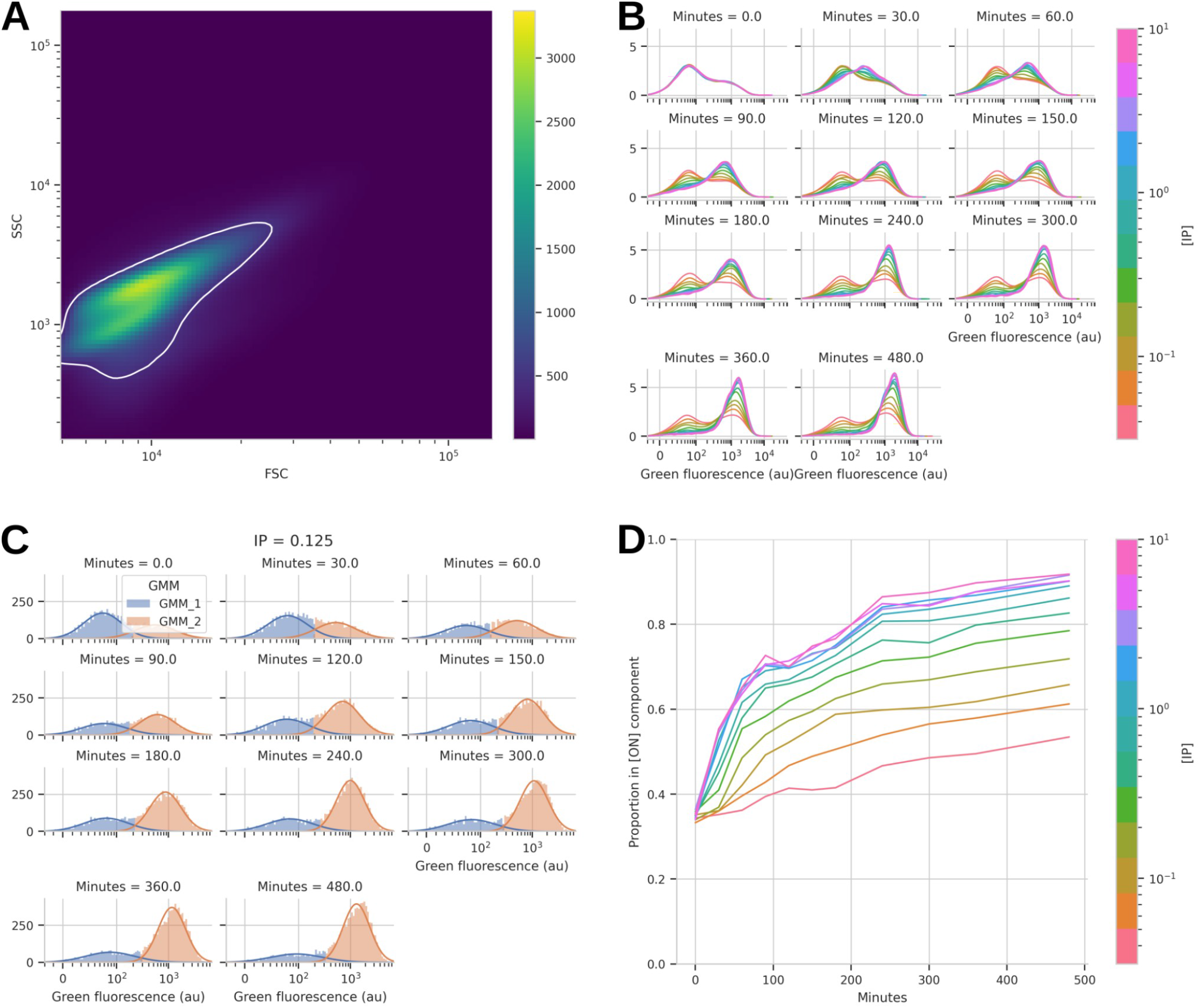
Cytoflow analysis of gene expression induced by a small molecule. (A) The population of single yeast cells was selected using a reproducible density gate instead of a hand-drawn gate. (B) A one-dimensional kernel density estimate shows that most experimental samples have bimodal distributions. (C) A one-dimensional, two-component Gaussian mixture model separates the two populations in each sample. The subplots are the various timepoints for the 0.125 μM concentration of the inducer; plots for other concentrations are similar. (D) A line graph of how the proportion of “ON” cells changes over both time and inducer concentration.

### Step 3: Visualize the fluorescence distributions

I used a kernel-density estimate – essentially a “smoothed” histogram – to visualize the data from the green fluorescence channel (Fig 5b) for events that are inside the density gate. Each sub-plot is the data from a single time point, and the colors on the plots represent the concentrations of inducer. Here, I noticed something very interesting: there is significant structure to this data. In contrast to a single unimodal distribution whose center increases in response to longer incubations or higher inducer concentrations, many of the distributions are bimodal! It’s almost as if there are two populations of cells – “off” and “on” – and the proportions of those two populations change over time.

### Step 4: Model distributions using a mixture model

If we hypothesize that each tube contains a mixture of two populations, an “off” population and an “on” population, then we might be able to apply a mixture model to better understand them. Though the data are not perfectly Gaussian, a Gaussian mixture model (GMM) may be useful to separate the two populations. Cytoflow includes a multidimensional GMM operation, which estimates the parameters for a GMM and assigns each event to the distribution with the highest posterior probability. Thus, I used the GMM operation to estimate a separate 1-dimensional, 2-component GMM for each FCS file (Fig 5c). This operation assigns each event to a component, but it also creates Statistics that record the estimated parameters from the GMM, including the center and spread of each component and the proportion of the overall data set that each component comprises.

### Step 5: Plot the summary data and draw conclusions

Because the GMM operation creates a Statistic containing the proportion of each component in the mixture, we can simply plot those proportions to get a better understanding of this strain’s induction dynamics and draw conclusions about its performance (Fig 5d). For example, it’s clear that the system “saturates” above about 1 μM of IP – additional IP does not change the way the strain behaves. It’s also clear that while the proportions of “off” and “on” cells change rapidly over the first 4 hours, they haven’t reached steady state even after 8 hours of induction. Finally, I think it’s very interesting that even the uninduced culture has a population of “on” cells, and that its proportion rises somewhat over the course of the experiment.

Together, these observations paint a much more comprehensive picture of the strain’s induction dynamics than a more traditional analysis. For example, if we simply compute the geometric mean fluorescence intensity (29) of each sample (Fig 6), we come away with a very different understanding: the saturation point is higher and the approach to “steady state” is much slower than with the GMM-based analysis.

**Figure 6.**
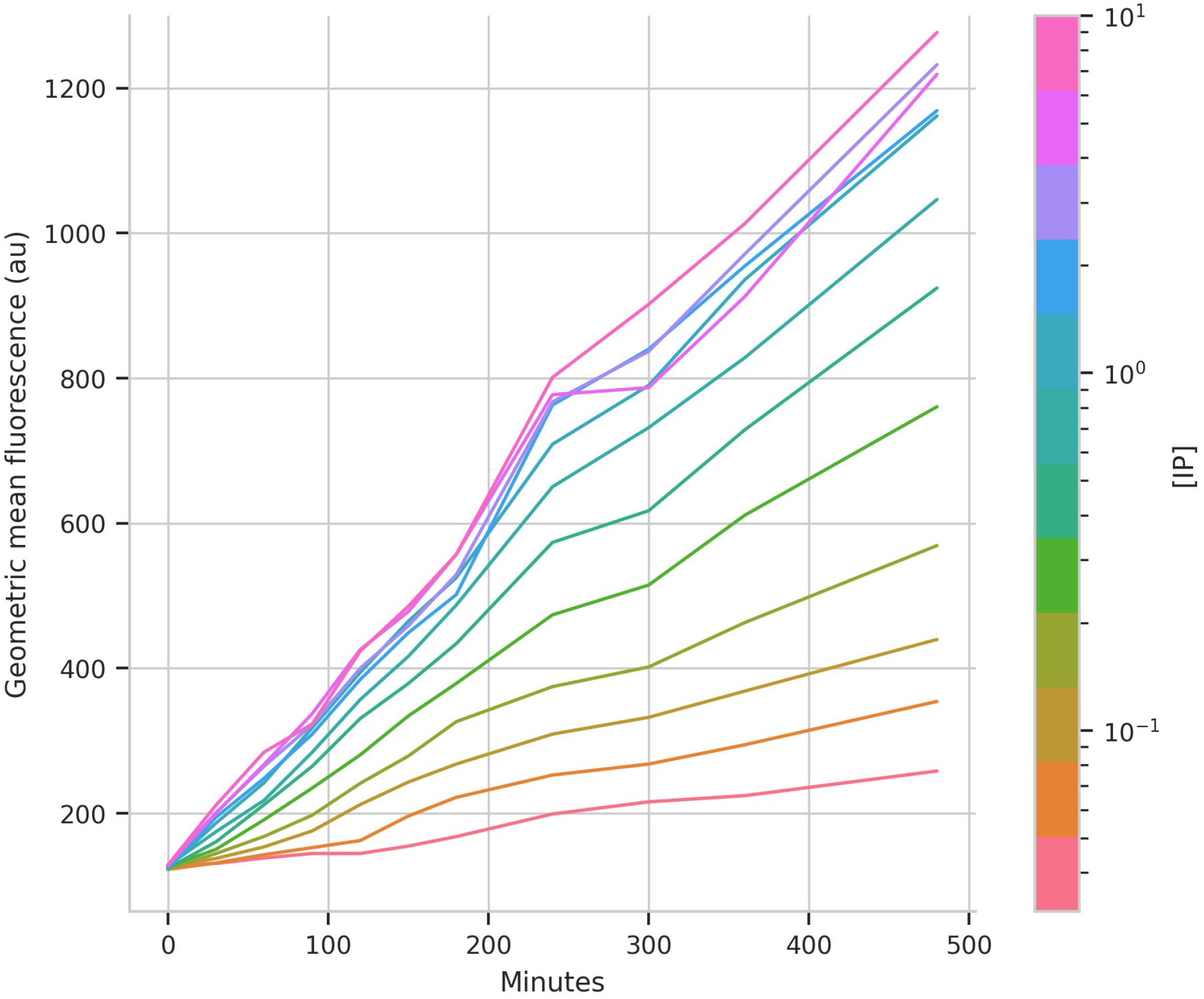
A more traditional analysis of gene induction leads to very different conclusions. Compared to Figure 5d, an analysis that simply computes the geometric mean of the cells’ fluorescence indicates that much more time, and a higher concentration of inducer, is necessary to fully induce this system.

## Availability and Future Directions

Cytoflow is an open-source project, and thus user input and contributions determine the direction of ongoing development. While Cytoflow’s functionality includes most of the “traditional” operations and visualizations, additional multidimensional classifiers, visualizations and analyses are of interest, particularly those developed specifically for analyzing flow cytometry data (23). Additionally, instruments based on new detection modalities such as spectral analysis, mass cytometry, and imaging cytometry may demand altogether new analyses. Bug reports, feature requests and patches are gratefully solicited! Code can be found at https://github.com/cytoflow/cytoflow, pre-built GUI binaries at https://cytoflow.github.io, and documentation at https://cytoflow.readthedocs.io.

## Acknowledgements

Initial development of Cytoflow was supported by United States National Science Foundation grant CNS-1446607 to Calin Belta and Douglas Densmore. My thanks to Nicholas DeLateur, Jacob Beal, Deepak Mishra, John Sexton, and Sebastian Castillo-Hair for critical feedback; the members of Ron Weiss’ lab and students in his courses for beta-testing; the Cytoflow users who have submitted bug reports and feature requests (and occasionally patches); and the developers of the scientific Python libraries on which Cytoflow relies, including **pandas** (30), **matplotlib** (31), **seaborn** (32), **numpy** (33), **scipy** (34), and others. Finally, I am grateful to the companies whose tools and resources support Cytoflow’s development, including Microsoft, Anaconda, Enthought, and Read the Docs, Inc.

